# Representations of semantic relations in the human cerebral cortex

**DOI:** 10.64898/2026.02.19.706815

**Authors:** Catherine Chen, Xue L. Gong, Fatma Deniz, Daniel L. Klein, Jack L. Gallant

## Abstract

An essential aspect of human cognition is the ability to explicitly think about semantic relations between concepts. Neuroimaging studies have found that individual concepts are encoded by distributed patterns of cortical activity, but it is unclear how representations of semantic relations compare to concept representations in the brain. Some theoretical models suggest that relation representations are embedded within concept representations, while others suggest that relation representations are independent of any specific concept pair. We designed a study to compare how semantic relations and concepts are encoded across the cerebral cortex. To characterize how relations are encoded across cortex, fMRI was used to record brain activity while six participants each answered over one thousand questions about different semantic relations. We find that representations of relations are distinct from their relata. Our results further suggest that relations and concepts are represented in similar cortical regions, and that, within these regions, each location is preferentially selective for specific relations. Overall, these results suggest that in the human cerebral cortex, relations and concepts have distinct representations that share a common functional organization.

## Introduction

To reason and communicate about the world, humans think both about individual concepts (such as *bicycle, wheel, vehicle*), and about how these concepts are semantically related to one another (for example, bicycle *has-part* wheel, bicycle *is-a* vehicle). The ability to think about individual concepts and their semantic relations is thought to be essential for human cognition. Note that throughout this paper we use the term “concept” to refer to a category (e.g., *bicycle*, *transportation*) that can be connected by a relation (e.g., *used-for*) (following Chaffin, 1989; Bejar et al., 1991, but see Reilly et al. 2024 for other ways in which this term has been used). This ability enables us to form generalizations, make inferences, and engage in analogical reasoning (Roger Chaffin 1989; Hofstadter and Sander 2013; Unger and Fisher 2021; Holyoak and Lu 2021; Bejar et al. 1991). Thus, understanding how the brain encodes semantic relations is crucial for understanding how it implements these flexible mental operations.

Neuroimaging studies have extensively mapped how concepts are encoded across cortex (Mitchell et al. 2008; Huth et al. 2016; Popham et al. 2021). These studies showed that concept representations are encoded in a network of brain regions distributed across the temporal, parietal, and prefrontal cortices, which together are often referred to as the *semantic system* (Binder et al. 2009; Huth et al. 2016). Each location is preferentially selective for certain concepts, forming a cortical map of concept representation that is largely consistent across individuals (Huth et al. 2016). However, the cortical map of relation representations remains unclear. Some studies have focused on taxonomic and thematic processing in the brain (Kalénine et al. 2009; Tsagkaridis et al. 2014; Mirman et al. 2017). Those studies used lesion analyses and neuroimaging contrasts, and found a potential neural dissociation in which ATL is engaged for taxonomic processing and TPC is engaged for thematic processing (but see Sachs et al. 2008; Jackson et al. 2015). A few studies have used representational similarity analysis (RSA) to compare brain representations of a few relation types, such as *is-a* and *has-part*, and found dissimilar brain responses for different relations (Chiang et al. 2021; Wang et al. 2021). These results suggest that different relations are represented at different locations in the brain. However, they do not provide a fine-grained functional map akin to what has been shown for concept representations. Such a map would allow us to directly inspect where each relation is represented, examine the functional organization of relation representations, and directly compare representations of relations to those of concepts.

Theoretical models suggest several possibilities for how brain representations of relations might compare to those of concepts. Some models suggest that relation representations are subordinate to concept representations. For example, relations are encoded in feature-comparison models as features of each concept (Smith et al. 1974), and in network models as edges between concept nodes (Collins and Quillian 1969). Other models suggest that relation representations are distinct from concept representations, such that relation representations are dynamically bound to different relata (Piantadosi et al. 2024). In semantic-relation networks, relations and concepts are feature vectors that are bound via synchronized firing (Hummel and Holyoak 1997; Doumas et al. 2004); in hyperdimensional computing models, relations and concepts are high-dimensional random vectors bound via element-wise vector addition (Kanerva 2009, 2010); in some vector space models, relations and concepts are embedding vectors bound via vector addition (Mikolov et al. 2013; Linzen 2016).

Behavioral evidence has shown that humans learn and use relations similarly to concepts (R. Chaffin and Herrmann 1984; Bejar et al. 1991; Roger Chaffin 1992; Popov et al. 2020). Based on this evidence, we hypothesized that relations may be represented in a similar way as concepts within the human cerebral cortex. Our hypothesis is consistent with the view that relation representations are distinct from representations of their relata. Our hypothesis additionally suggests that representations of relations and concepts have a similar functional organization: this would mean that relations are encoded in similar cortical regions as concepts, and each location within these regions is preferentially selective for certain relations. To test our hypothesis, we designed a study to map and compare functional brain representations of relations and of concepts.

## Results

In this study, six participants each performed a relation-verification experiment. In this experiment, each participant answered over 1000 questions involving eight different relations. In each trial of the experiment, participants answered questions about one of eight relations. The eight relations included six different semantic relations and two different non-semantic relations. The six semantic relations were: *is-a* (e.g., *bicycle-vehicle*), *found-at* (e.g., *bicycle-garage*), *has-part* (e.g., *bicycle-wheel*), *symbol-of* (e.g., *bicycle-sustainability*), *used-for* (e.g., *bicycle-transportation*), and *made-of* (e.g., *bicycle-aluminum*). These six semantic relations were chosen for two reasons. First, they are hypothesized to reflect how humans classify the relations between concepts. For this reason, they have been used in several prior studies of semantic relations (Bejar et al. 1991; Jurgens et al. 2012). Second, relations can be applied to a wide range of objects. This allowed us to use a consistent set of objects for trials probing different relations, which ensured that any observed similarities or differences in brain representations across relations would reflect differences in the relation type, rather than differences in the concepts involved in the trials probing each relation. The objects used in this experiment were all human-made artifacts (a full list is in Table S1). To distinguish representations of semantic versus non-semantic relations, the experiment also included trials of two non-semantic wordform relations: *alphabetically-before* (e.g., bicycle-stone), and *wordform-match* (e.g., bicycle-bicycle). Throughout the rest of this text we refer to semantic relations simply as <u>relations</u>, and the two non-semantic wordform relations as <u>wordform relations</u>.

We used voxelwise encoding models to map functional brain representations of each relation. One feature space reflected the six relations. This feature space was a six-dimensional binary vector that described the relation type for each trial. To account for additional features that might confound the results of our study, we constructed sixteen additional feature spaces. These additional feature spaces account for the two wordform relations, the lexical semantic content of the presented words, visual aspects of the stimuli, motor reactions, participant reaction times, and participant accuracy. Banded ridge regression was used to jointly estimate model weights that predict the BOLD activity in each voxel from all sixteen feature spaces (Nunez-Elizalde et al. 2019). The estimated model weights were used to predict BOLD activity on a held-out test set. The held-out test set comprised new instances of each relation that were not used for model estimation. The product measure was used to quantify the contribution *R~^2^* of each feature space to the total prediction accuracy (Hoffman 1960; Pratt 1987; Dupré la Tour et al. 2022). Model weights were estimated and evaluated separately in each voxel and for each individual participant.

We first test whether relation representations are independent from specific concept pairs. In our experiment, the held-out test set contained new concept pairs that were not included in the train set. Thus, if relation representations are independent from specific concept pairs, then relation model weights estimated from the train set should accurately predict brain responses in the held-out test set, which contains different concept pairs from the train set. Alternatively, if relation representations are tied to specific concept pairs, then the relation model weights will not generalize to the held-out test set.

Figure 2a shows the prediction accuracy (√*R~^2^_relation_*) of the relation model weights on the held-out test set. Results are shown separately for each individual participant, and at the group-level. For each individual and at the group-level, brain responses are significantly well-predicted throughout bilateral temporal, parietal, and prefrontal cortices (p<.05 by a one-sided permutation test, after a Benjamini-Hochberg correction for multiple comparisons (Benjamini and Hochberg 1995)). These results show that relation model weights generalize to new instances of each relation, and so suggest that the representations of these relations are independent of specific concept pairs.

**Figure 1:**
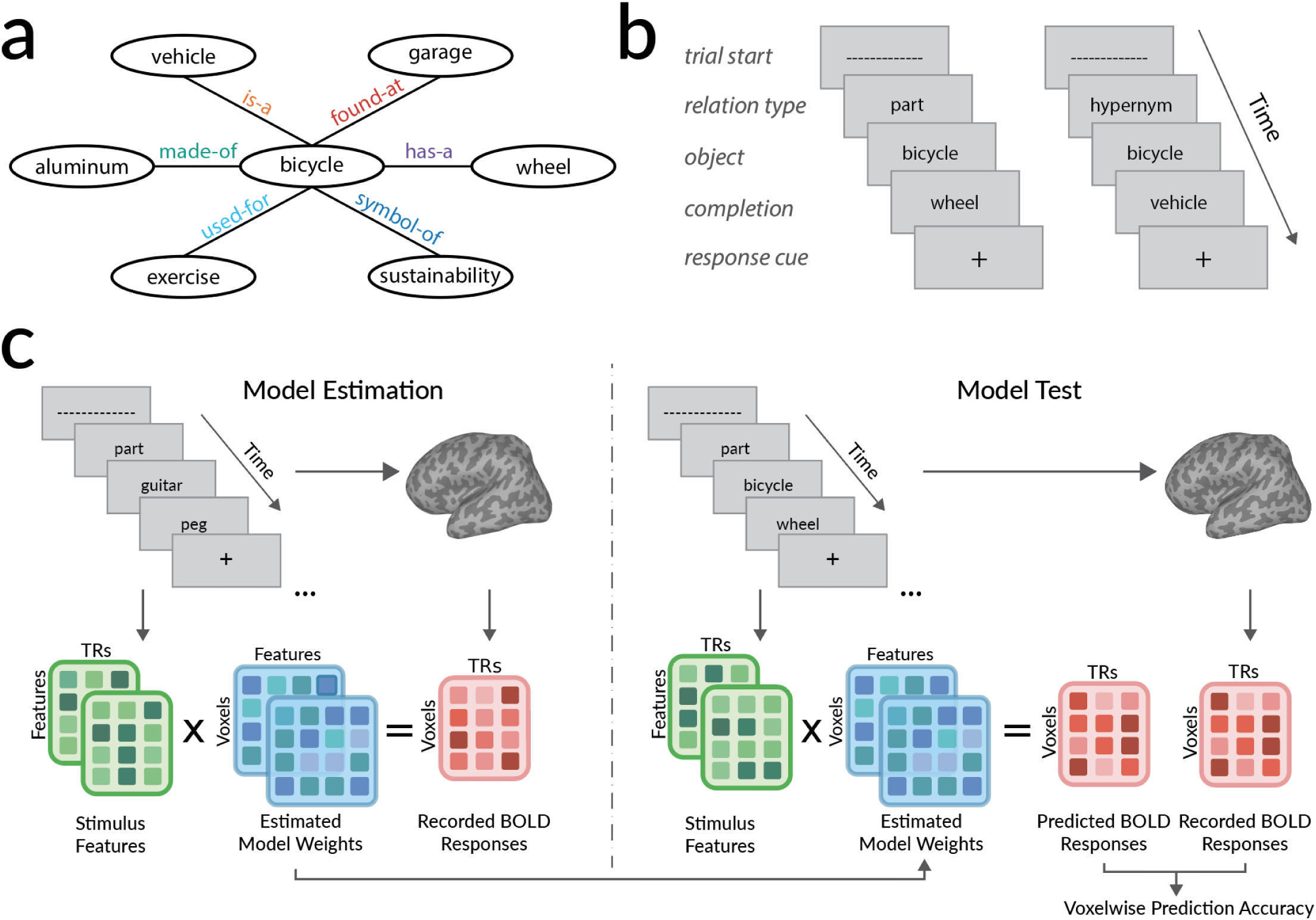
**a**. Examples of the six relations in our experiment. Examples are shown for the object “bicycle”. **b**. Experiment paradigm. Participants each performed over 1000 trials of an event-related relation-verification experiment while fMRI was used to record BOLD activity. Two example trials are shown. In each trial, displayed words indicated a relation (e.g., “part”), an object (e.g., “bicycle”), and a potential completion (e.g., “wheel”). Participants were instructed to press a button to indicate whether the presented instance forms a valid relation. **c**. Modeling framework. For each relation a binary feature space was constructed to describe the times at which participants performed trials of that relation. Voxelwise modeling was used to estimate a separate FIR encoding model for each feature space, voxel, and participant. Estimated model weights describe how each relation modulates BOLD activity separately in each voxel and for each participant. Estimated model weights were used to predict BOLD activity to a held-out dataset that was not used for model estimation. The held-out dataset used different instances of each relation from the train dataset. Prediction accuracy was quantified as the coefficient of determination *R*^2^ between predicted and recorded BOLD activity in the held-out dataset.

**Figure 2:**
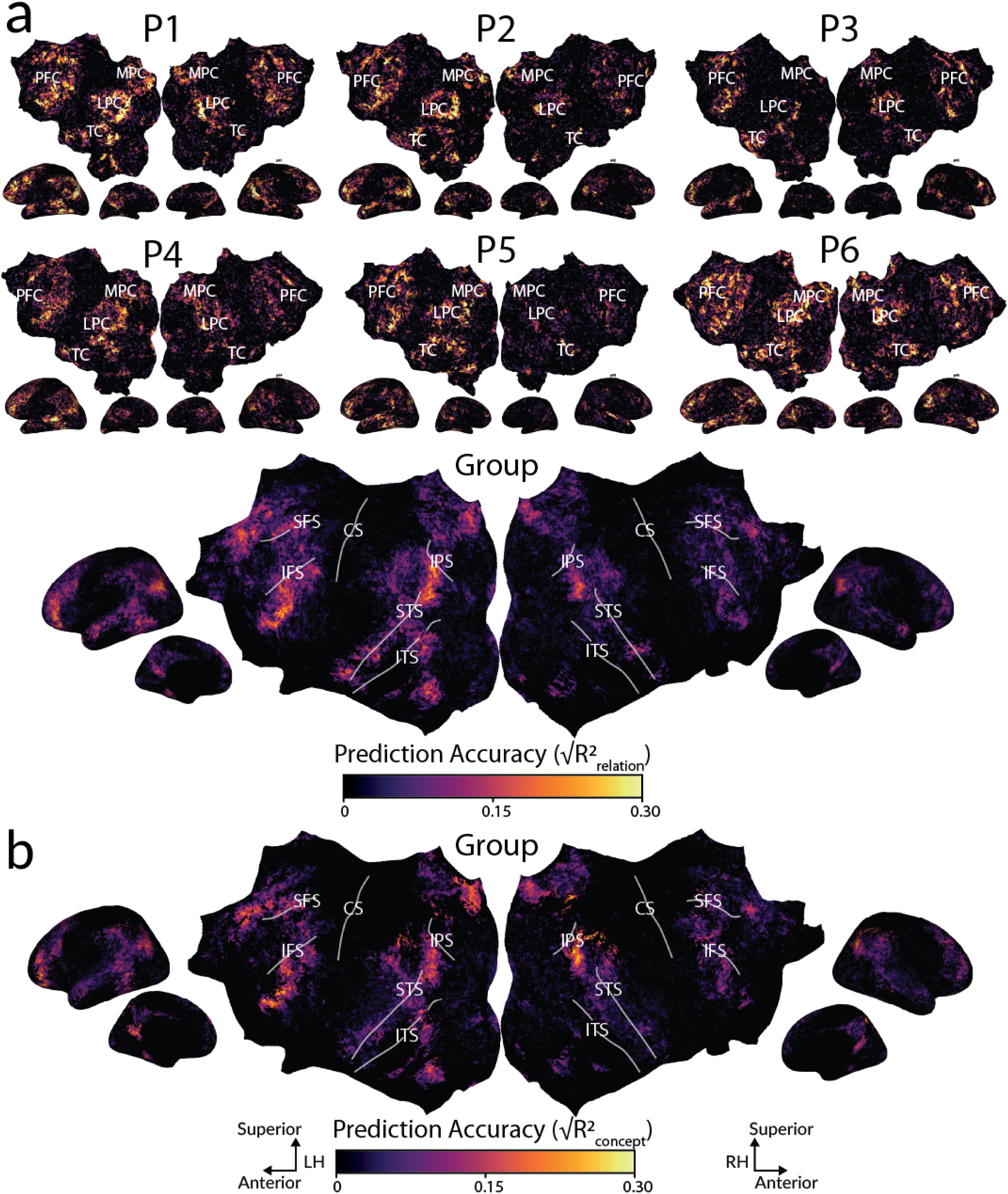
**a.** Relation representations. The relation-verification experiment was used to estimate relation representations across the cerebral cortex. To determine whether relations have their own representations, we tested whether the estimated relation model weights could accurately predict BOLD activity for held-out instances of each relation. For each participant, prediction accuracy of the relation model weights is shown in the participant’s native brain space. Group-level accuracies were obtained by projecting voxelwise accuracies for each participant to a standard template space (fsaverage7; (Fischl et al. 1999)) and then taking the average across participants for each vertex of the template space. Group-level results are shown on the flattened cortical surface of the template space. Each voxel or vertex is colored according to the prediction accuracy of the relation model weights. Voxels that are not significantly well-predicted are shown in black (one-sided p<.05, FDR corrected). For each participant and at the group-level, prediction accuracy is highest in the bilateral temporal, parietal, and prefrontal cortices. These results show that relation representations are consistent across different concept pairs. **b.** Concept representations. Prediction accuracy of the lexical semantic model weights is shown at the group-level, in the same format as (a). Results for each individual participant are consistent with the group and are shown in Figure S1. Prediction accuracy is highest in the bilateral temporal, parietal, and prefrontal cortices. These results show that relations and concepts are encoded in the same set of brain regions. (STS=superior temporal sulcus, ITS=inferior temporal sulcus, IPS=intraparietal sulcus, SFS=superior frontal sulcus, IFS=inferior frontal sulcus).

The results in Figure 2a show that relations are encoded throughout the temporal, parietal, and prefrontal cortices. Previous studies have used lexical semantic features to model concept representations in the brain, and reported that these brain regions also encode concepts (Mitchell et al. 2008; Binder et al. 2009; Huth et al. 2016; Deniz et al. 2019). Those results suggest that similar brain regions encode both relations and concepts. To verify this inference we examined the prediction accuracy of the lexical semantic model in each of the six participants in our study.

Figure 2b shows the prediction accuracy of the lexical semantic model weights (√*R~^2^_lexical semantic_*) on the held-out test set, at the group-level. (Results for each individual participant are shown in Figure S1, and are consistent with the group.) Prediction accuracy is highest within bilateral temporal, parietal, and prefrontal cortices. Voxels throughout these regions are significantly well-predicted (p<.05 by a one-sided FDR-corrected permutation test). Furthermore, voxelwise prediction accuracy for each participant is positively correlated between the relation model and the concept model (r=0.38, 0.34, 0.28, 0.28, 0.25, 0.39 for P1-P6 respectively; p<.001 FDR corrected permutation test, for each individual participant). Thus, relations and concepts appear to be encoded in similar brain regions.

Having established that relations and concepts are encoded in similar cortical regions, we next examined whether within these regions each location is preferentially selective for certain relations. Prior work has extensively mapped the functional organization of concept representations. That work has shown that each voxel is preferentially selective for certain concepts, and that the cortical organization of concept selectivity is largely consistent across individuals (Huth et al. 2016; Deniz et al. 2019; Popham et al. 2021; Chen, Gong, et al. 2024). To determine whether relations follow a similar organizational principle, we tested whether each voxel that encodes relations is preferentially selective for certain relations over others. Voxelwise selectivity for each relation is reflected by the relation model weights: if a voxel is selective for a certain relation, this means that brain activity in the voxel is relatively higher when the participant thinks about that specific relation, and the corresponding model weight will be high. To determine whether each voxel is preferentially selective for certain relations, or broadly selective for all relations, we tested whether selectivity for one relation was proportionally much higher than selectivity for other relations. In this test, we are agnostic to which specific relation (e.g., *is-a* vs. *has-part*) a voxel is selective for; we are interested in whether selectivity for any one of the relations is much higher than selectivity for the other relations. Therefore for this analysis, for each voxel we sort the relations in order from highest to lowest selectivity, and refer to the ordered relations simply as *R*_0_ (highest selectivity) through *R*_5_ (lowest selectivity). For each voxel, we then examine the proportional selectivity for each *R_i_*, which was determined by dividing the selectivity for *R_i_* by the total selectivity for all six relations.

Proportional selectivity is shown in Figure 3a. Results are plotted separately for each participant. Because the purpose of this analysis is to determine whether voxels that encode relations are preferentially selective for specific relations, voxels that do not encode relations (√*R~^2^_relation_* < 0.1) were excluded from Figure 3. Visual inspection suggests that for all six participants, most voxels are preferentially selective for one relation. To ensure that this pattern is not due to chance, a jackknife procedure was used to identify voxels in which selectivity is significantly greater for one relation than for any of the other five relations (Abdi and Williams 2022). Figure 3b shows the set of voxels with significant selectivity, plotted for each participant separately. For every one of the six participants, more than 90% of voxels that are well-predicted by the relation feature space have only one single preferred relation (P1: 96%, P2: 96%, P3: 92%, P4: 92%, P5: 94%, P6: 96%). This suggests that, like for concepts, most voxels are preferentially selective for certain relations.

**Figure 3:**
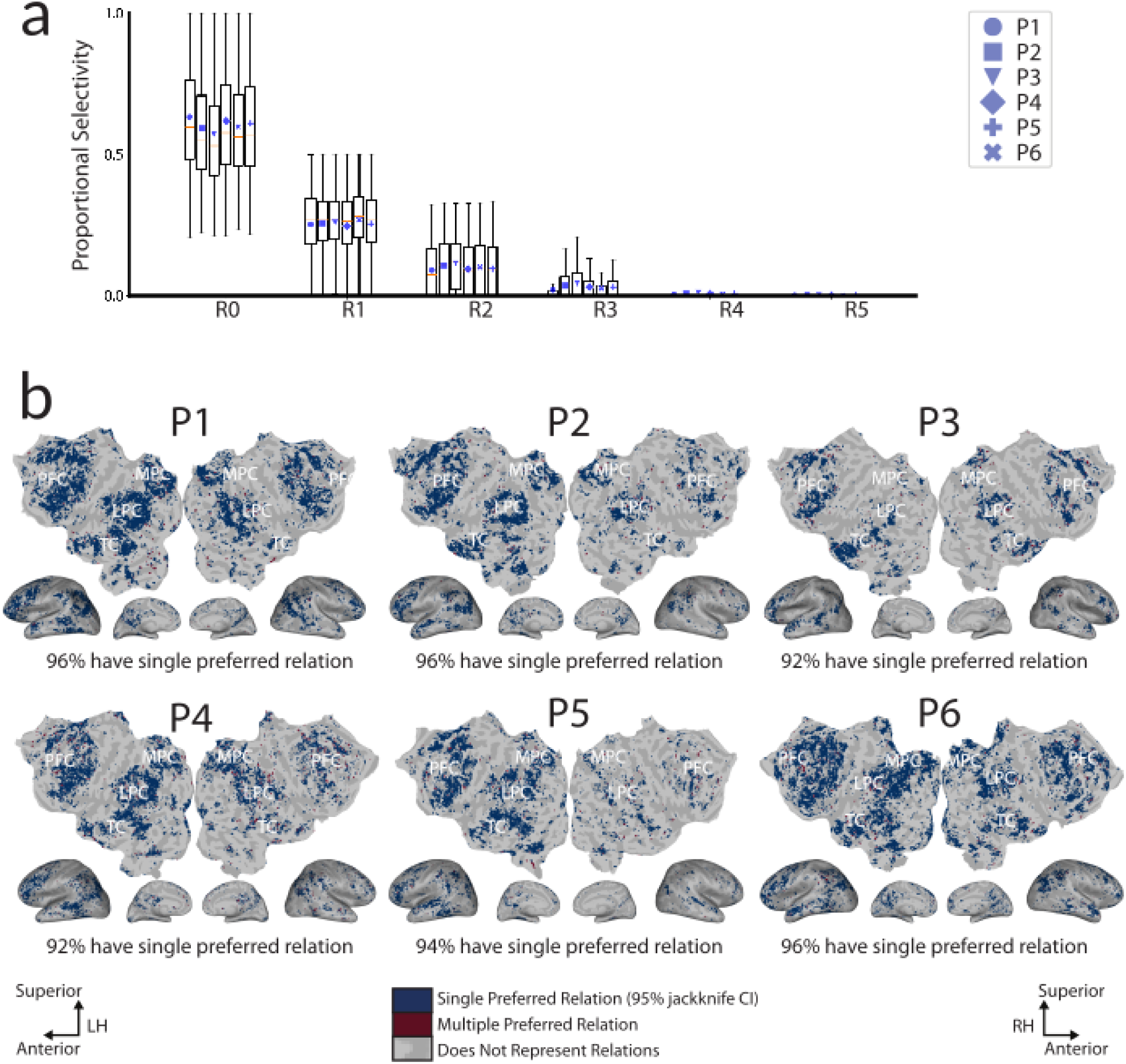
To determine whether each voxel is preferentially selective for certain relations or is broadly selective for all relations, we examined each voxel’s proportional selectivity for each relation. For this test we are agnostic to the identity of the relation (e.g., *is-a* vs. *has-part*) for which a voxel is most selective; therefore we sort the relations in order from highest to lowest selectivity, and refer to the ordered relations simply as *R*_0_ (highest selectivity) through *R*_5_ (lowest selectivity). **a**. Proportional selectivity for *R*_0_ through *R*_5_. Each boxplot reflects the distribution over voxels in one participant. Across participants, most voxels are preferentially selective for one relation. **b.** A jackknife procedure was used to test the statistical significance of each voxel’s preference for a single relation. Results are shown for each participant on the flattened surface of the native brain space. Voxels shown in blue prefer a single relation. Voxels shown in red prefer multiple relations. Voxels shown in grey do not represent relations. Very few voxels are shown in red. In every one of the six participants, over 90% of voxels prefer a single relation (P1: 96%, P2: 96%, P3: 92%, P4: 92%, P5: 94%, P6: 96%). These results suggest that most voxels that encode relations are preferentially selective for certain relations.

Figure 3 established that most voxels are preferentially selective for one of the six relations in our experiment, but ignored the identity of the preferred relation of each voxel. Thus, it is unclear whether the cortical organization of relation representations is, like that of concept representations, consistent across participants. To evaluate whether this is the case, the preferred relation of each voxel was first projected from the participant’s native brain space to the common fsaverage7 template space. Figure 4 shows preferred relations for each vertex of the template space. (Results in native participant space are consistent with results in the template space, and are shown in Figure S2). Results are shown at the group-level and in each individual participant. Visual inspection suggests that the cortical organization of preferred relations is largely consistent across participants. In lateral parietal cortex (LPC) *found-at* (red), *has-part* (purple), *is-a* (orange), and *symbol-of* (blue) are arranged in a ring of patches located around the temporoparietal junction. In medial parietal cortex (MPC) *found-at*, *is-a*, and *symbol-of* are located in patches in, superior to, and anterior to retrosplenial cortex. In temporal cortex (TC) *found-at*, *made-of*, and *symbol-of* are located anterior to parahippocampal place area, superior to parahippocampal place area, and along the superior temporal sulcus. In prefrontal cortex (PFC) *symbol-of* is located superior to superior frontal sulcus and inferior to inferior frontal sulcus, and *found-at* is located along superior frontal sulcus. To quantify this result, for each pair of participants we measured the percentage of template vertices that have the same preferred relation between the two participants. (Note that the match between participants can never reach 100%, because of measurement noise and anatomical and functional variation between participants.) For all 15 pairs of participants, the percentage of vertices with the same preferred relation is above 48%, and this value is significantly greater than chance (p<.001 by a permutation test, FDR corrected). These results suggest that relation representations are organized in cortical patterns that are generally consistent across individuals.

**Figure 4:**
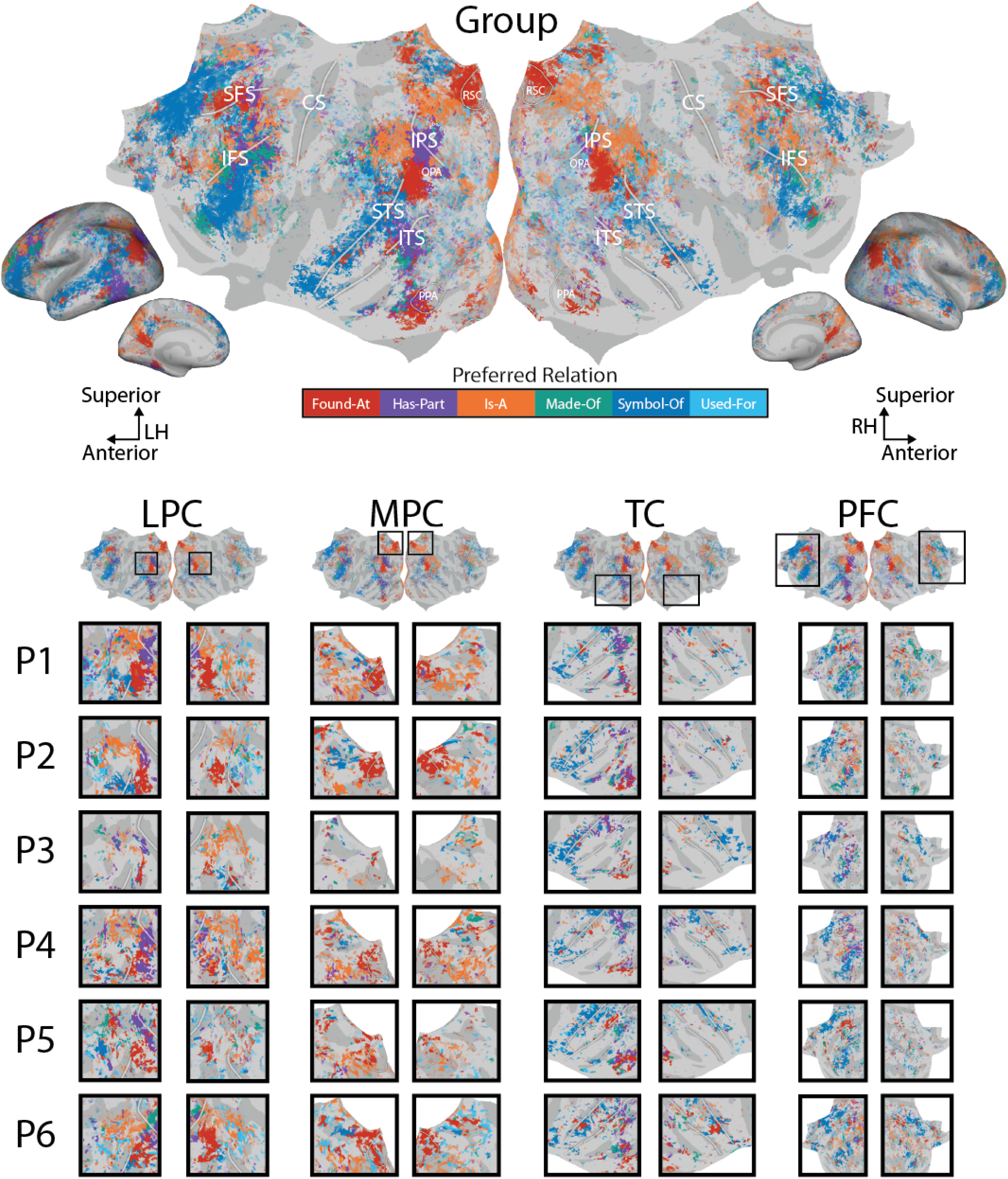
We examined whether the cortical distribution of preferred relations is consistent across participants. Results are shown at the group-level and for each individual participant. To facilitate comparisons across participants, all results are shown on the flattened cortical surface of the fsaverage7 template space. The color of each vertex reflects the preferred relation. The opacity of each vertex reflects the prediction accuracy of the relation model weights. Vertices with multiple preferred relations and those that do not encode relations are shown in grey. Visual inspection suggests that the cortical organization of preferred relations is consistent across participants. In lateral parietal cortex (LPC) *found-at* (red), *has-part* (purple), *is-a* (orange), and *symbol-of* (blue) are arranged in a ring of patches located around the temporoparietal junction. In medial parietal cortex (MPC) *found-at*, *is-a*, and *symbol-of* are located in patches in, superior to, and anterior to retrosplenial cortex (RSC). In temporal cortex (TC) *found-at*, *made-of*, and *symbol-of* are located anterior to parahippocampal place area (PPA), superior to PPA, and along the STS. In prefrontal cortex (PFC) *symbol-of* is located superior to SFS and inferior to IFS, and *found-at* is located along SFS. At least 48% of vertices have consistent preferred relations between every pair of participants (p<.001, by a permutation test). These results suggest that the cortical organization of relation representations is generally consistent across participants.

The previous results suggest that relations are encoded independently from the specific concepts they connect, and that the functional representation of relations is similar to that of concepts: relations are encoded in similar cortical regions as concepts, and each location within these regions is preferentially selective for certain relations.

One potential confound in our study is that the estimated relation representations may merely reflect the influence of the specific words involved in trials of each relation. While trials of all relations involve the same set of core objects (Table S1), each relation connects these objects to different words. For example, trials of the *made-of* relation include words such as *glass* (cup *made-of* glass) and *aluminum* (bicycle *made-of* aluminum) while trials of the *found-at* relation include words such as *kitchen* (cup *found-at* kitchen) and *garage* (bicycle *found-at* garage). To avoid this potential confound, the lexical semantic features of each stimulus word were included as a nuisance feature space during model estimation. The relation model weights nevertheless accurately predict brain responses, which suggests that our relation model reflects brain representations of the relations themselves.

To further validate that the estimated relation representations are not confounded by word semantics, we inspected voxelwise concept selectivity. For brevity, we will hereafter refer to these words that differ across trials for different relations as “relation-specific words”. If there is indeed a confound and estimated relation representations merely reflect the semantics of the relation-specific words, then the lexical semantic weights of a voxel that prefers a certain relation should be close to the corresponding set of relation-specific words. For example, a voxel that prefers the *made-of* relation should be highly selective for words that appeared in *made-of* trials, such as *glass* and *aluminum*. Alternatively, if the voxels that prefer a certain relation are selective for concepts beyond the corresponding set of relation-specific words, then it would suggest that the estimated relation representations are *not* confounded by word semantics.

To perform this validation, for each voxel we computed the mean Pearson correlation between the lexical semantic weights of that voxel and the embeddings of the relation-specific words for the preferred relation of the voxel. Figure 5 shows the distribution over voxels of these correlations. Results are separated by preferred relation, and are shown for each participant separately. Across relations and participants, voxels exhibit low correlations. For each participant, at least 25% of voxels exhibit a negative correlation (P1: 33%, P2: 34%, P3: 33%, P4: 36%, P5: 33%, P6: 28%). This result shows that the voxels that prefer a certain relation are not necessarily selective for the corresponding set of relation-specific words. This suggests that the estimated relation representations are not confounded by word semantics.

**Figure 5:**
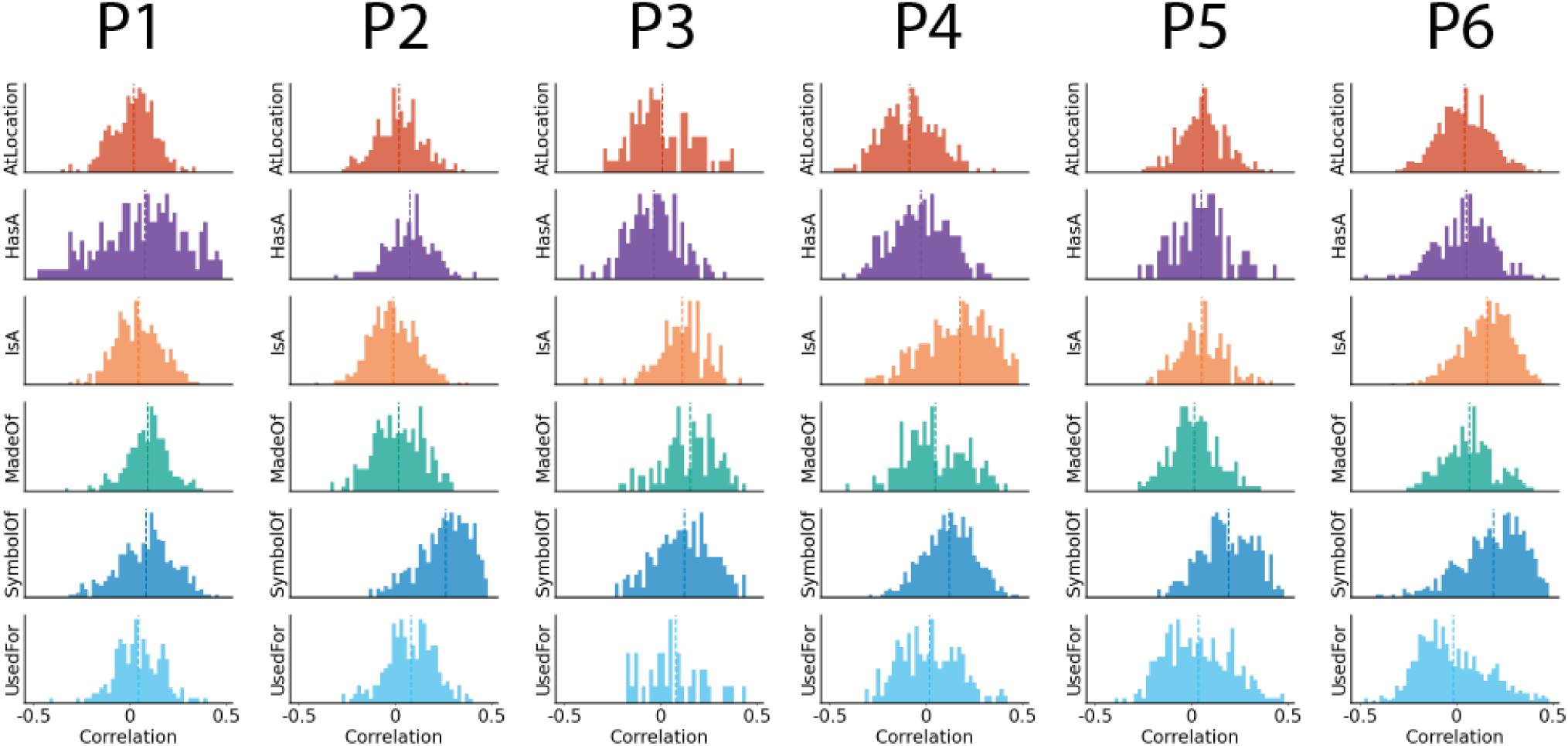
To test whether estimated relation representations are confounded by word semantics, we tested whether a voxel that is selective for a certain relation is selective for the words that appeared in trials of that relation (the corresponding “relation-specific words”). If there is indeed a confound and estimated relation representations merely reflect the semantics of the relation-specific words, then the lexical semantic weights should closely align with the embeddings of the relation-specific words. For each voxel, the Pearson correlation was computed between the lexical semantic weights of the voxel and the embeddings of the corresponding relation-specific words. Histograms show the distribution of correlations over voxels, separately for each preferred relation and participant. Across relations and participants, voxels exhibit low correlations. For each participant, at least 25% of voxels exhibit a negative correlation (P1: 33%, P2: 34%, P3: 33%, P4: 36%, P5: 33%, P6: 28%). These results suggest that the estimated relation representations are not confounded by word semantics.

## Discussion

We hypothesized that relations and concepts may have distinct representations that share a common functional organization. The results presented here provide three lines of evidence in support of this hypothesis. First, relations are encoded independently from specific concept pairs. Second, relations are encoded in similar cortical regions as concepts. Third, like for concepts, each location within these regions is preferentially selective for certain relations. These results generalize across participants, including the two held-out participants (P5 and P6).

Prior neuroimaging studies of relation representations in the brain did not directly compare representations of relations and concepts. Some studies examined the geometry of concept representations in the brain and found that this geometry linearly encodes information about semantic relations (Zhang et al. 2020; Wu et al. 2022; Ferrante et al. 2025). For example, the brain representations of *teacher*, *chalk*, and *mechanic* can be added and subtracted (*mechanic* + *chalk* - *teacher*) to approximate the brain representation of *wrench*. However, this latent structure may not be what the brain draws upon during relation processing. Thus, we designed a study to specifically target relation processing and directly map representations of relations. Our results are consistent with prior studies which found that brain responses are similar across different instances of the same relation (Wang et al. 2021; Chiang et al. 2021). Moreover, we directly modelled how each relation is encoded in voxel responses and thereby showed that relation representations have a similar functional organization as concept representations.

In designing this study we sought to avoid any potential confound that might occur if our estimates of relation selectivity partly reflected the influence of the specific words involved in trials of each relation. (For example, trials of the *made-of* relation include words such as *glass* and *wood* while trials of the *found-at* relation include words such as *kitchen* and *garage*.) To mitigate against this potential confound, the same set of objects was used across trials of all six relations, and the lexical semantic features of each stimulus word were included as a nuisance feature space during model estimation. Thus, the high prediction accuracy of the relation model weights suggests that our relation model reflects brain representations of the relations themselves, rather than the words involved in trials of each relation. Furthermore, if the relation model were confounded by word semantics, then we would expect that for each relation, the voxels that prefer that relation will also have concept selectivities that are closest to the words involved in trials of that relation. However, Figure 5 shows that this is not the case. Thus, it is unlikely that the estimated relation representations are confounded by the semantics of the words involved in trials of each relation.

Another potential concern is that the statistical significance of preferred relations may be an artifact of regularization hyperparameters. Because the selectivity of each relation depends on the magnitude of the model weights, regularization hyperparameters influence the estimates of relation selectivity. However, in our models all six relations are part of the same feature space, which means that for each voxel, the same regularization hyperparameter is used for all six relations. For each voxel, the significance test for preferred relations only considers relative differences between selectivity for each relation. Thus, the statistical significance of preferred relations is not confounded by regularization hyperparameters.

One limitation of this study is that it focuses solely on the cerebral cortex. Relations may be encoded differently in other parts of the brain. For example, relations may be encoded in the hippocampus, which is thought to play a key role in supporting relational knowledge, and to index to distributed representations in the neocortex (Teyler and DiScenna 1986; Cohen et al. 1999; Eichenbaum 2000; Garvert et al. 2017). Relations may also be encoded in the cerebellum, which has been implicated in tasks involving semantic integration and semantic associations (e.g., (Argyropoulos and Muggleton 2013; Gatti et al. 2020)). The methods developed here could be used to investigate the representation of relations in the cerebellum and in deep brain structures.

To avoid potential confounds from the objects involved in each relation, our study controlled the objects and relations in the stimuli. The methods developed here could easily be adapted to examine additional concept categories and relation types. In particular, our method could be used to compare relations involving abstract and concrete concepts, examine whether relation representations vary under different task settings such as analogy problems or unstated relations, and test hypotheses about similarities between relations (Roger Chaffin 1989).

Our study identified relation-selective patches throughout many brain regions. We speculate that these patches may form a hierarchical architecture for relation processing. Relation-selective patches in occipital, lateral parietal, and temporal regions are broadly consistent with sensorimotor processing of the modalities associated with certain relations. For example, we found selectivity for “found-at” in regions typically associated with scene processing (RSC, PPA, and OPA); we found selectivity for “has-part” in lateral occipitotemporal regions associated with visual object processing; we found selectivity for “used-for” in regions such as left posterior STS and SMG typically associated with action processing (Kuhnke et al. 2023). Relation patches identified in lateral parietal cortex and temporal cortex may correspond to sensorimotor and verbally mediated representations of relations; patches near prefrontal cortex may support goal-directed behaviors related to relations, such as determining which relation to emphasize in different situations; patches in medial parietal cortex may support episodic memory, such as retrieving and storing memories mediated by certain relations. This hierarchical architecture would be consistent with prior proposals that semantic memory may form a hierarchical architecture in which modality-specific processing occurs around sensorimotor processing regions, top-down selection of semantic information occurs in prefrontal cortex, and encoding and retrieval of episodic memory occurs in regions such as the posterior cingulate gyrus and precuneus (Binder and Desai 2011).

The results presented here can inform theoretical models of how relations are encoded in the human brain. While some models suggest that relations are encoded as edges between concept nodes or as features contained within concept representations (Collins and Quillian 1969; Smith et al. 1974), others suggest that relations have their own representations that are dynamically bound to concept pairs (Kanerva 2010; Gentner and Forbus 2011; Holyoak et al. 2022; Doumas and Hummel 2012). Our results show that relation representations generalize across different pairs of concepts, and support the view that relation representations are distinct from any specific pair of connected concepts. Furthermore, we find similarities in how concepts and relations are represented in the brain: both are represented in the same general set of brain regions, and voxels within these regions show preferential selectivity for specific concepts and specific relations. These results lend support to the view that relations and concepts have distinct but analogous representations in the brain (Hummel and Holyoak 1997; Kanerva 2010; Holyoak et al. 2022).

## METHODS

### Experimental Design

The relation-verification experiment followed a trial-based design. Each trial consisted of three words. The first word indicated a relation (“hypernym” to indicate “is-a”, “part” to indicate “has-part”, “material” to indicate “made-of”, “purpose” to indicate “used-for”, “environment” to indicate “found-at”, or “idea” to indicate “symbol-of”; participants were instructed on the correspondence between each word and relation before the experiment). The second word indicated an object (e.g., “bicycle”). The third word indicated a potential completion (e.g., “vehicle”). The experiment included 1496 separate trials for each participant. The trials were evenly divided across eight relations (six semantic relations and two non-semantic wordform relations). The same set of 65 common objects was used across trials of each relation. Half of the trials contained true examples (e.g., “hypernym-bicycle-vehicle”) and half contained false examples (e.g., “hypernym-bicycle-clothing”).

At the start of each trial a dotted line was presented for 0.4 seconds. Then a rapid serial visual presentation (RSVP) procedure (Forster 1970) was used to present the three words one-by-one at the center of the screen. Presentation duration was determined by word length. Each word was presented for a base length of 0.3 seconds, and 0.01 seconds were added for each letter of the word. Each triple of words was followed by an answer period that lasted between 1 and 3 seconds. Answer period durations were jittered to ensure that trials were not time-locked to TR onsets. During the answer period participants were asked to press a button to indicate whether the triple of words formed a valid instance of a relation, answering either “True” or “False”. During the answer period no words were present on the screen.

Trials were presented across 11 unique runs. Each run contained 136 trials. The order of trials was randomized. Relation types, objects, and correct answers (true or false) were balanced across runs. One of the 11 runs was used as a test run. This run contained instances of each relation that were not used in any of the ten train runs. The test run was performed twice by each participant (once in each session). The two repeats of the test run were used to get an estimate of the noise ceiling (Sahani and Linden 2002; Hsu et al. 2004; Schoppe et al. 2016).

Participant reaction time was quantified as the time elapsed between the onset of the answer cue and the participants’ response. The distribution of participant reaction time is shown in Figure S3. For each trial, the participant’s response was considered to be correct if their response was the same as the most common response among the other five participants. Note that this is a conservative estimate of participant accuracy, because for some trials the answers are subjective (e.g., *bicycle*-*symbol of*-*sustainability*). The distribution of participant accuracy is shown in Figure S4.

Functional MRI data were collected from each participant during two 2-hour scanning sessions that were performed on different days. Each scanning session consisted of six ten-minute long runs (five were train runs and one was the test run).

### Participants

Functional data were collected from six naive participants between the ages of 25-29 (5 female, 1 male). All participants were right handed according to the Edinburgh handedness inventory (Oldfield 1971) (+90, +85, +75, +90, +100, +55 for P1-P6; laterality quotient of −100: entirely left-handed, +100: entirely right-handed). The entire experiment was piloted extensively on the first author of this study. Those pilot data are not included in this publication because the author overlearned the stimuli during experimental design and piloting.

To ensure generalization across participants the entire analysis was performed in each individual participant and consistency was measured between each participant and the group. Furthermore, before performing any analyses two of the six participants (P5, P6) were designated as held-out participants (Popham et al. 2021). Data for these two participants were not analyzed until the entire analysis and modeling pipeline was finalized. Results were consistent between the two held-out participants and the other four participants.

### fMRI Data Acquisition

Whole-brain MRI data were collected on a 3T Siemens TIM trio scanner located in the Henry J. Wheeler Brain Imaging Center at UC Berkeley. A 32-channel Siemens volume coil was used. Functional scans for participants P1, P2, P3, P5, and P6 were collected with a T2*-weighted gradient-echo EPI with repetition time (TR)=2.0045s, echo time (TE)=35ms, flip angle=74°, voxel size=2.24×2.24×4.1 mm (slice thickness=3.5mm), matrix size=100×100, and field of view=224×224 mm. Thirty axial slices were prescribed to cover the entire cortex and were scanned in interleaved order. A custom-modified bipolar water excitation radiofrequency (RF) pulse was used to prevent contamination from fat signals. The functional scans for participant P4 were collected with a sequence with multiband acceleration factor 3, repetition time (TR)=1.156s, echo time (TE)=34ms, flip angle=62°, voxel size=2.5×2.5×2.5 mm (slice thickness=2.5mm), matrix size=84×84, and field of view=210×210 mm. Anatomical data were collected with a T1-weighted multi-echo MP-RAGE sequence on the same 3T scanner. The results for participant P4 were consistent with the other five participants, suggesting that the difference in functional scanning parameters did not affect the results of this study.

To stabilize head motion during scanning sessions and to ensure that each participant was centered within the receive coil on each session, participants wore a personalized head case that precisely fit the shape of each participant’s head (Gao 2015; Power et al. 2019).

### fMRI data pre-processing

The FMRIB Linear Image Registration Tool (FLIRT) from FSL 5.0 was used to motion-correct each functional run (Jenkinson et al. 2002). All volumes in the run were averaged across time to obtain a high quality template volume. FLIRT was used to automatically align the template volume for each run to the overall template, which was chosen to be the temporal average of the first functional run for each participant. The cross-run transformation matrix was then concatenated to the motion-correction transformation matrices obtained with MCFLIRT, and the concatenated transformation was used to resample the original data directly into the overall template space. Noise from motion, respiratory, and cardiac signals were removed with a component-based detrending method (CompCor; (Behzadi et al. 2007)). Responses of each run were z-scored separately. During z-scoring, for each voxel separately the mean response across time was subtracted and the remaining response was scaled to have unit variance. Before data analysis, 10 TRs from the beginning and 10 TRs at the end of each run were discarded in order to account for the 10 seconds of silence at the beginning and end of each scan, and for non-stationarity in brain responses at the beginning and end of each scan.

### Cortical surface reconstruction and visualization

FreeSurfer software was used to generate cortical surface meshes from the T1-weighted anatomical scans (Fischl et al. 1999). Before surface reconstruction, anatomical surface segmentations were carefully hand-checked and corrected with Blender software and pycortex (Gao et al. 2015; Community 2018). Relaxation cuts were made into the surface of each hemisphere. Blender and pycortex were used to remove the surface crossing the corpus callosum. The calcarine sulcus cut was made at the horizontal meridian in V1 using retinotopic maps data as a guide.

Pycortex was used to align functional images to the cortical surface (Gao et al. 2015). The line-nearest scheme in pycortex was used to project functional data onto the surface for visualization and analysis. This projection scheme samples the functional data at 32 evenly spaced intervals between the inner (white matter) and outer (pial) surfaces of the cortex and then averages together the samples. Samples are taken using nearest-neighbor interpolation, wherein each sample is given the value of its enclosing voxel.

### Statistical Analyses

Voxelwise modeling (VM) was used to model the recorded BOLD activity (Naselaris et al. 2011; Nunez-Elizalde et al. 2019; Dupré la Tour et al. 2022, 2025). In the VM framework, stimulus and task parameters are first transformed nonlinearly into feature spaces that reflect variables of interest. Each feature space describes an aspect of the experiment that is hypothesized to be encoded in brain responses. Linearized, regularized regression is used to estimate a separate encoding model for each voxel and feature space. The model weights describe how each feature space modulates the BOLD activity of each voxel. A held-out dataset that was not used to estimate model weights is used to evaluate prediction accuracy and to determine the statistical significance of prediction accuracy.

All model fitting and analysis was performed with custom software written in Python, making heavy use of NumPy (Harris et al. 2020), SciPy (Virtanen et al. 2020), and Matplotlib (Hunter 2007).

#### Stimulus Feature Spaces

A six-dimensional binary feature space was constructed to reflect the timing of trials for each relation. Each dimension of this feature space corresponds to one of the six relations in the experiment, and reflects the onset time and duration of trials involving that specific relation. An analogous two-dimensional binary feature space was constructed to reflect the timing of trials for each wordform relation. Fifteen nuisance feature spaces were constructed to reflect potentially confounding variables: visual-spatial and motion features (motion energy) (Adelson and Bergen 1985; Watson and Ahumada 1985; Nishimoto et al. 2011), the letters in each word, word length, standard deviation of word length within each TR, the mean word length per TR, the change in mean word length across TRs, word rate, the length of each trial, the length of each question, the participant reaction time, whether each TR is part of a question or answer period, the correct answer to the trial, the participant’s answer, the accuracy of the participant’s answer, and the lexical semantics of each word. The lexical semantics of each word was operationalized as the word2vec embedding of each word that was shown on the screen.

#### Stimulus Feature Preprocessing

Before voxelwise modeling, each stimulus feature was truncated, downsampled, mean-centered, and delayed. Data for the first 10 TRs and the last 10 TRs of each scan were truncated to account for the 10 seconds of silence at the beginning and end of each scan, and for non-stationarity in brain responses at the beginning and end of each scan. An anti-aliasing, 3-lobe Lanczos filter with cut-off frequency set to the fMRI Nyquist rate (0.25 Hz) was used to resample the stimulus features to match the sampling rate of the fMRI recordings. Then the stimulus features were each mean-centered. Lastly, finite impulse response (FIR) temporal filters were used to delay the features in order to model the hemodynamic response function of each voxel. The FIR filters were implemented by concatenating feature vectors that had been delayed by 2, 4, 6, and 8 seconds (following e.g. Huth et al. 2016; Deniz et al. 2019; Chen, Dupré la Tour, et al. 2024). A separate FIR filter was fit for each feature, participant, and voxel.

For one participant (P1) the presented stimuli inadvertently contained some specific trials that were repeated in both the train and test runs. To remove this overlap, the train stimulus features were cleaned by setting the relation stimulus feature spaces to 0 for any trial that was also included in the test stimuli for participant P1. This train feature cleaning procedure ensured that the estimated model weights did not reflect information from trials that appeared in the test set. Results were consistent between P1 and the other five participants, suggesting that this train feature cleaning procedure did not affect our results.

#### Regularization hyperparameter selection

Five-fold cross-validation was used to find the optimal regularization hyperparameters for each feature space and each voxel. Hyperparameter candidates were chosen with a random search procedure (Bergstra and Bengio 2012): 1000 normalized hyperparameter candidates were randomly sampled from a Dirichlet distribution, and were then scaled by 10 log-spaced values ranging from 10^−5^ to 10^5^. The regularization hyperparameters for each feature space and voxel were selected as the hyperparameters that produced the minimum squared error (L2) loss between the predicted voxel responses and the recorded voxel responses (*argmin*_ℎ*yperparameters*_ ||*y*^ − *y*||_2_^2^). The voxels with the lowest 20% of the cross-validated L2 loss were selected for refinement in the second stage. The second stage consisted of 1000 iterations of hyperparameter gradient descent (Bengio 2000). Regularization hyperparameters were chosen separately for each participant and voxel. This hyperparameter search was performed with the Himalaya Python package (Dupré la Tour et al. 2022).

#### Model estimation and evaluation

The selected regularization hyperparameters were used to estimate model weights that map from the relation feature space to voxel BOLD activity. Model weights were estimated separately for each voxel and participant. For each relation, the model weights for the corresponding dimension of the relation feature space reflect voxelwise selectivity for that relation.

The test set was not used to select hyperparameters or to estimate regression weights. The prediction accuracy *R*^2^ of the feature spaces was computed per voxel as the coefficient of determination between the predicted and recorded voxel responses on the test set.

The statistical significance of prediction accuracy was computed by a permutation test with 1000 iterations (Deniz et al. 2019; Chen, Dupré la Tour, et al. 2024; Reddy and Wehbe 2020; Tang et al. 2024). In each permutation iteration, the timecourse of voxel responses in the test dataset was permuted by blockwise shuffling (shuffling was performed in blocks of 10 TRs in order to preserve autocorrelations in voxel response), and then the prediction accuracy *R*^2^ was computed between the permuted and predicted response for each voxel separately. The distribution of test accuracies over permutation iterations was used as a null distribution to compute the p-value of prediction accuracy for each voxel.

#### Group-level prediction accuracy

Group-level prediction accuracy was computed by first computing prediction accuracy for each participant in the participant’s native brain space, and then projecting individual participant results into a standard template space (fsaverage7; (Fischl et al. 1999)). Average prediction accuracy across six participants was computed for each fsAverage vertex.

#### Voxelwise Relation Selectivity

If a voxel is *selective* for a particular relation, it means that the voxel will be more highly activated when a participant thinks about that relation. Voxelwise relation selectivity is reflected by the model weights of the six-dimensional relation feature space. The relation feature space is a binary feature space where each dimension of this feature space corresponds to one of the six relations in the experiment. If a voxel is selective for a particular relation, then the voxel will be more highly activated during trials of that relation as compared to other points in the experiment, and the model weights for the corresponding dimension will be high.

### Voxelwise Preferred Relations

A jackknife procedure was used to determine the statistical significance of voxel preference for a specific relation. For each voxel the six relations were sorted from highest to lowest selectivity. The ordered relations were referred to as *R*_0_ (highest selectivity relation) through *R*_5_ (lowest selectivity relation). A jackknife procedure was used to estimate confidence intervals around the difference between selectivity for *R*_0_ and for each of *R_i_* ∈ {*R*_1_, *R*_2_, *R*_3_, *R*_4_, *R*_5_} (Abdi and Williams 2022). The jackknife procedure is a leave-one-out resampling approach in which items are iteratively dropped to construct partial estimates of the parameter of interest. All samples are used to construct a whole-sample estimate of the parameter of interest, and the partial estimates are compared to the whole-sample estimate in order to estimate the standard error of the parameter. This standard error can then be used to construct confidence intervals. We use the jackknife procedure with a leave-one-run-out approach. Each of the 10 train runs was left out in turn, and the remaining 9 runs were used to re-fit the voxelwise encoding model and construct a partial estimate of voxelwise selectivity for each of the relations. These partial estimates were used to estimate 95% confidence intervals around *S_r_*_0_ − *S_ri_* for *i* ∈ {1, 2, 3, 4, 5}. A strictly positive confidence interval suggests that the observed higher selectivity for *R*_0_ as compared to *R_i_* is unlikely to be due to chance. A voxel was considered to significantly prefer a specific relation if all five confidence intervals were strictly positive.

For each voxel, the relation with the highest selectivity was defined as the *preferred relation* of that voxel. Voxels that were not well-predicted by the relation feature space (√*R~^2^_relation_* < 0.1) were considered to have no preferred relation.

To compare the organization of preferred relations across participants, voxelwise preferred relations for each participant were projected to the template space. For each pair of participants, the set of vertices that are well-predicted by the relation feature space in both participants was identified. Then, consistency of preferred relations was measured as the percentage of these vertices that have the same preferred relation between the two participants.

## Acknowledgements

This work was supported by the following grant awards: National Science Foundation (NSF Nat-1912373); German Federal Ministry of Education and Research (BMBF 01GQ1906); Berliner ChancengleichheitsProgramm (BCP) to FD; a National Science Foundation GRFP (DGE 1752814) to CC; an IBM PhD Fellowship to CC; and by internal UC Berkeley funds. We thank Jen Holmberg, Chris Kymn, Kevin Lin, Anwar Nunez-Elizalde, Jerry Tang, Christine Tseng, and Alicia Zeng for their feedback on the study and manuscript.

## Supplementary Information

**Figure S1.**
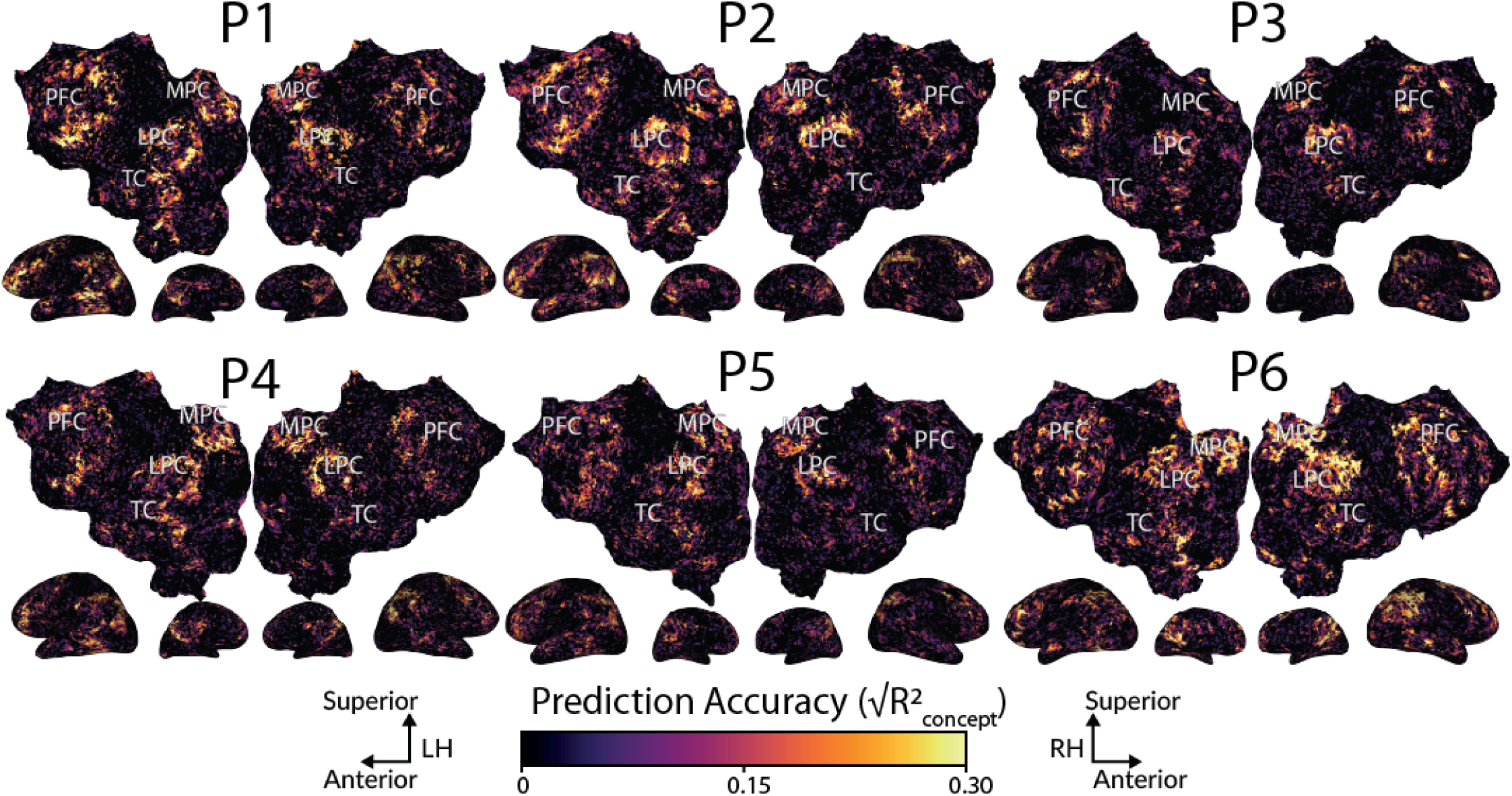
Cortical distribution of concept representations, for each participant in the participant’s native brain space. The format is the same as in Figure 2b. Relations and concepts are represented in the same set of brain regions.

**Figure S2:**
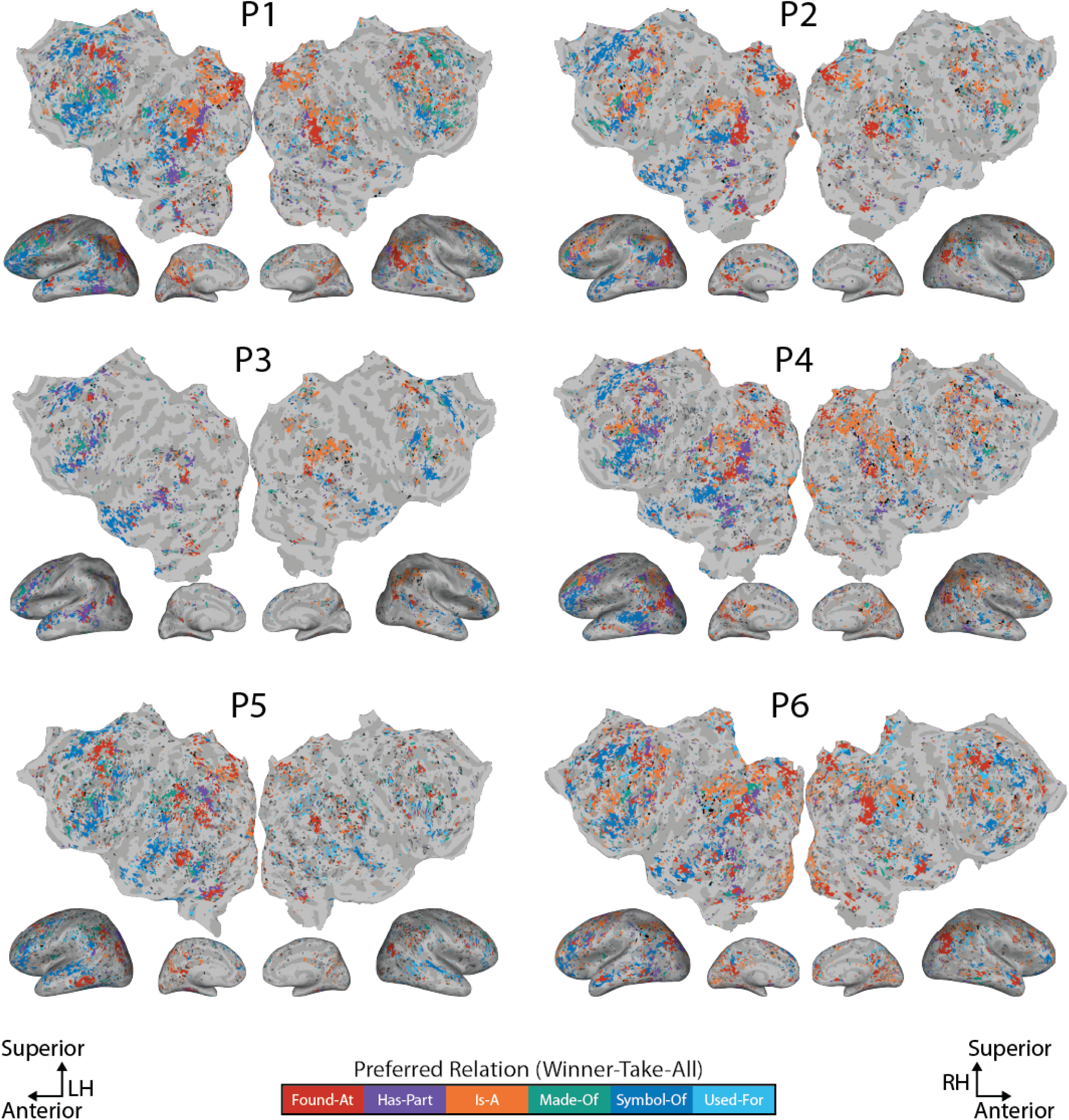
Cortical organization of preferred relations, for each participant in the participant’s native brain space. The format is similar to Figure 4. The cortical organization of preferred relations appears to be consistent across participants.

**Figure S3:**
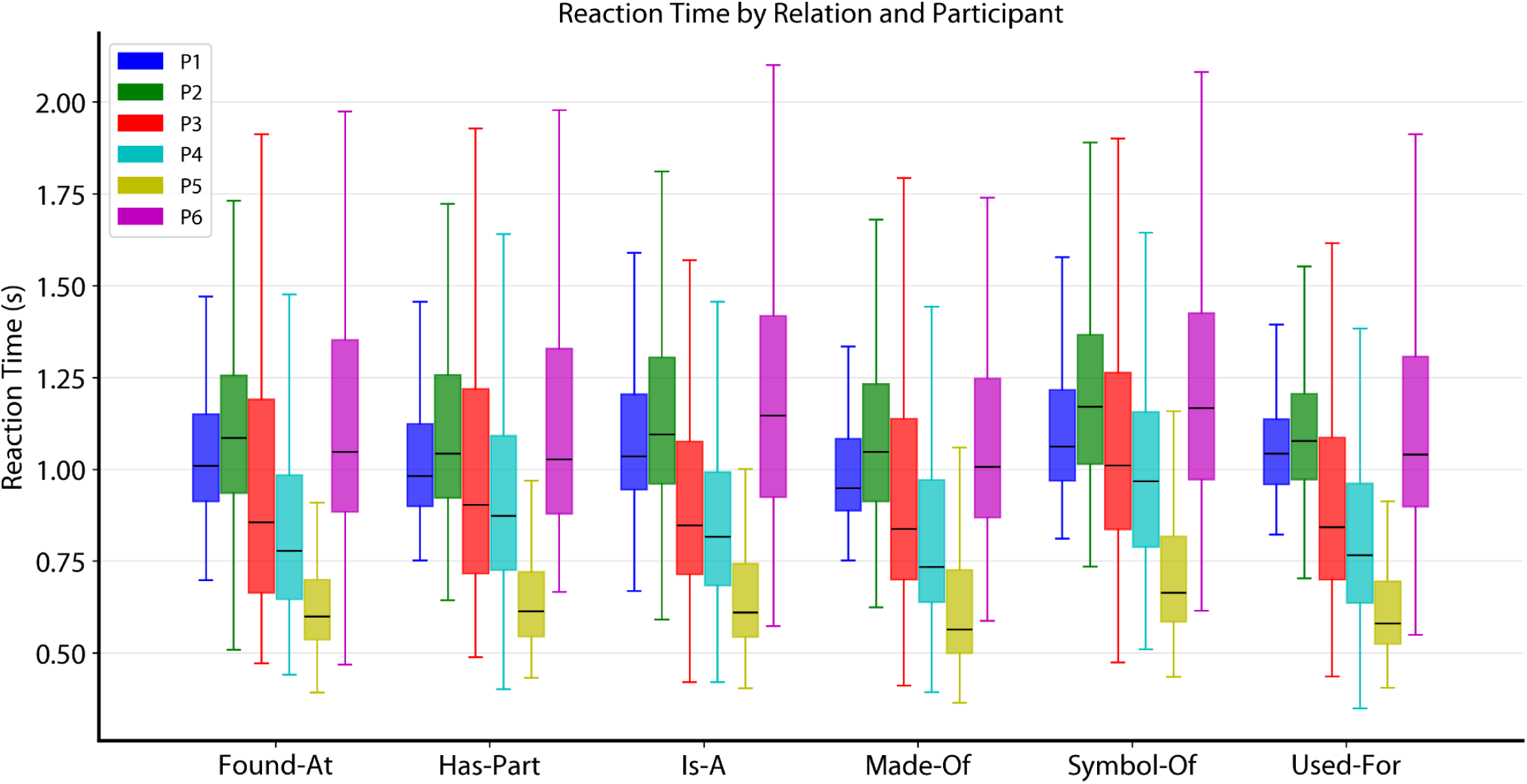
Mean reaction time for each relation and participant. The reaction time of each trial was defined as the time between when the question was shown and the participant’s response. In each boxplot, the line indicates the median over trials. The box indicates the first and third quartile over trials. The whiskers indicate the farthest data points within 1.5x of the inter-quartile range from the boxes. The median reaction time primarily varies by participant.

**Figure S4.**
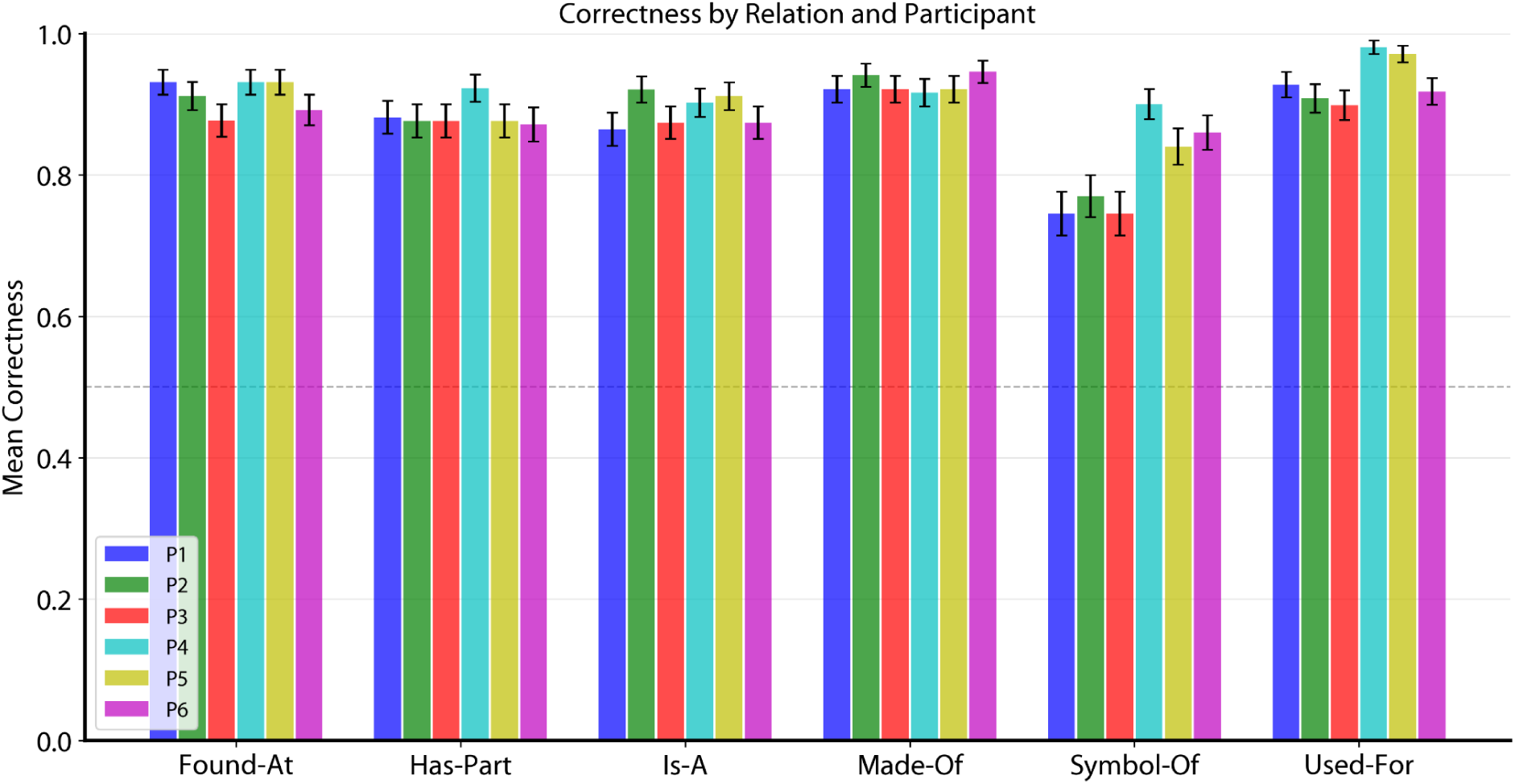
Mean accuracy for each relation and participant. For each participant, the accuracy of each trial was defined based on whether the participant agreed with the mode of the other five participants. Bars indicate the mean accuracy over trials, for each participant and relation separately. Error bars show the standard error over trials. For most relations, mean accuracy is above 80%. Accuracy of “Symbol-of” is lower than for the other relations, possibly reflecting the relatively subjective nature of this relation.

**Table S1:** List of object words used in the experiment.

|  |  |  |  |  |  |  |
|---|---|---|---|---|---|---|
| airplane | camera | gun | microscope | ship | table | viola |
| armchair | candle | hammer | motorcycle | shoe | telescope | violin |
| axe | car | harp | necklace | shovel | toilet | wallet |
| backpack | cellphone | jacket | oven | sled | tractor | wrench |
| bed | couch | kettle | pan | spatula | train | wristwatch |
| bicycle | dress | knife | pencil | stove | truck |  |
| boat | flashlight | magazine | piano | submarine | trumpet |  |
| book | flute | mailbox | purse | sweater | typewriter |  |
| briefcase | fork | metronome | rake | swimsuit | ukelele |  |
| cabinet | guitar | microphone | screwdriver | sword | van |  |

